# pyMAP: a Python package for small and large scale analysis of Illumina 450k methylation platform

**DOI:** 10.1101/078048

**Authors:** Amin Mahpour

**Affiliations:** Cancer Genetics Department, Roswell Park Cancer Institute, Buffalo NY 14263, USA

## Abstract

PyMAP is a native python module for analysis of 450k methylation platform and is freely available for public use. The package can be easily deployed to cloud platforms that support python scripting language for large-scale methylation studies. By implementing fast parsing functionality, this module can be used to analyze large scale methylation datasets. Additionally, command-line executables shipped with the module can be used to perform common analysis tasks on personal computers.

Availability and implementation: PyMAP is implemented in Python and the source code is available under GPL v2 license from http://aminmahpour.github.io/PyMAP/.

## Introduction

Illumina Infinium 450k BeadChip platform has been critical for methylation analysis of human cells. Indeed, this platform has been largely used in The Cancer Genome Atlas (TCGA) to assess the methylation status of many different type of tumors [1]. Since methylation study of genomes in normal and neoplastic conditions are usually performed in large scale clinical studies, a powerful yet simple tool is needed to organize, parse and analyze the resulting methylation datasets. This tool should also implemented to be extensible. Here, we describe Python package for Methylation Analysis of Probes(PyMAP) to analyze data generated from the 450k platform. This package will significantly reduce the hassle associated with parsing and analyzing the methylation profile of large scale projects. Importantly, as a free and open-sourced project, this package could be further modified to fulfill the need of special research and clinical projects.

## Implementation

PyMAP is implemented in pure and fast python code and does not require any prerequisite package to parse and analyze methylation data. In contrast, existing R packages are dependent on other packages and therefore are very difficult to deploy on personal and lab workstations [2]. This package fully supports Python 3.4 and the latest version 3.5 and is also backward compatible with legacy python version of 2.7.

Multi-tasking of certain processes is also supported to reduce the processing time associated with large studies. Therefore, we think efficient use of processing power and memory usage contributes to better user experience and reasonable requirements for computational resources.

PyMAP contains two essential sub-modules of Core and Annotations along with others for a typical analysis pipeline. Core module implements parse functions that can efficiently process single, or in batch form, multiple methylation data files that are exported from Illumina GenomeStudio Methylation software. Our performance testing showed that the package can parse 60 data samples in less than 20 seconds with extremely low memory footprint on a Unix workstation. In addition, Annotation module fully supports Illumina probes and associated information. We have also added the ability to filter probes that are associated with Single Nucleotide Polymorphisms. Additionally, this module allows customized filtering of probes for regions of interests in genome. In other words, probes that are associated with a certain location, gene, feature or CpG site in the genome can be retrieved easily and in combination with core module, methylation data can be analyzed.

The advantage of implementing this package in pure python is manifold. Python allows broad scalability and fast deployment to workstation and cloud services [3]. Indeed, well known cloud processing services such as Galaxy bioinformatics platform and Amazon Web Services already support python and therefore setting up large scale analysis is far more convenient in these powerful platforms. Additionally, the code base also allows efficient consumption of the module in other modules and standalone python applications.

## Features

Streamlined extraction of probes that are associated with a genomic feature and associated methylation beta values are extremely important and we implemented this functionality in a way that is intuitive. The extracted probes can be used to fetch the methylation values of samples and generate methylation profile associated with a certain genomic feature. Fast parsing of methylation data through Core module stores all methylation information necessary for analysis of large methylation dataset. For preprocessing, PyMAP can remove SNP associated probes which is a common process for differential methylation analysis in human samples [4]. PyMAP also supports differential methylation analysis of probes using Student t-test statistics method. However, due to the modular pipeline, other methods for differential analysis can also implemented to the current pipeline.

In addition, PyMAP supports extraction of the methylation data into formatted BED files that can be visualized in UCSC genome browser or any other genome browser that support genomics BED files as specified on UCSC genome browser website. Additionally, by implementing convenient functions, large scale generation of BED files for multiple sample is also feasible. In addition, these exported BED files can be used with Circos program for genome-wide visualization of the methylation data. Also, we have created fully documented and easy to use executable command-line scripts to perform common analysis and designed specifically for use of researchers who are not familiar with the python script.

We also have added a plotting module to plot the methylation values of desired probes from selected samples. This is particularly useful if the user would like to visually inspect the methylation values of a filtered probe list associated with a feature or genomic region.

## Documentation

PyMAP is developed with added emphasis on public availability and contains an easy to follow documentation. The code is also highly documented to encourage customization by end users. The project full documentation is available at http://pymap.readthedocs.org.

## Future directions

PyMAP, as a script module, provides a complete pipeline for analysis of a large scale methylation dataset. However, we have intentions to develop an interactive Graphical User Interface(GUI) in near future that consumes full PyMAP functionality in a graphically rich environment. The standalone application would further simplify the analysis and visualization of the methylation data for scientists.

